# Stroke prevents exercise-induced gains in bone microstructure but not composition in mice

**DOI:** 10.1101/708388

**Authors:** Nicholas J. Hanne, Andrew J. Steward, Marci R. Sessions, Hannah L. Thornburg, Huaxin Sheng, Jacqueline H. Cole

**Author notes:** **Corresponding Author:** Jacqueline H. Cole Joint Department of Biomedical Engineering University of North Carolina andNorth Carolina State University 911 Oval Drive Campus Box 7115, Raleigh, NC 27695-7115.

## Abstract

Ischemic stroke induces rapid loss in bone mineral density that is up to 13 times greater than during normal aging, leading to a markedly increased risk of fracture. Little is known about skeletal changes following stroke beyond density loss. In this study we use a mild-moderate middle cerebral artery occlusion model to determine the effects of ischemic stroke without bedrest on bone microstructure, dynamic bone formation, and tissue composition. Twenty-seven 12-week-old male C57Bl/6J mice received either a stroke or sham surgery and then either received daily treadmill exercise or remained sedentary for four weeks. All mice were ambulatory immediately following stroke, and limb coordination during treadmill exercise was unaffected by stroke, indicating similar mechanical loading across limbs for both stroke and sham groups. Stroke did not directly detriment microstructure, but exercise only stimulated adaptation in the sham group, not the stroke group, with increased bone volume fraction and trabecular thickness in the sham distal femoral metaphysis. Stroke differentially decreased cortical area in the affected limb relative to the unaffected limb of the distal femoral metaphysis, as well as endosteal bone formation rate in the affected tibial diaphysis. Although exercise failed to improve bone microstructure following stroke, exercise increased mineral-to-matrix content in stroke but not sham. Together, these results show that stroke inhibits exercise-induced changes to femoral microstructure but not tibial composition, even without changes to gait. Similarly, affected-unaffected limb differences in cortical bone structure and bone formation rate in ambulatory mice show that stroke affects bone health even without bedrest.

## INTRODUCTION

Stroke is the leading cause of long-term disability in the United States – approximately 7.0 million Americans have had a stroke, and nearly 4% of the population are projected to have a stroke by 2040 [1]. In addition to cognitive and motor impairments, stroke also severely affects skeletal health, particularly in the paretic limbs, by reducing bone mineral density (BMD) up to 13% per year compared to 1% for healthy aging individuals over 60 years of age [2–7]. Reduced BMD and increased susceptibility to falling lead to a 15% fracture incidence within 5 years following stroke and a 47% increased risk of fracture compared to age- and sex-matched controls [8,9]. Traditionally, BMD loss following stroke has been attributed to paresis and bedrest. However, in a study examining BMD a year following severe stroke in completely bed-ridden patients, stroke patients still lost more BMD in their paretic limbs, suggesting stroke impacts skeletal health beyond the effects of mechanical unloading [6]. Aside from BMD loss and increased fracture risk, little is known about the effects of stroke on bone health. Understanding more about how stroke affects other bone measures, particularly at the tissue and cellular levels, is a critical first step for identifying mechanisms underlying bone fragility post-stroke and mitigating bone loss in these patients.

Middle cerebral artery occlusion (MCAo) is a well-established technique to induce ischemic stroke in rodents. The MCAo-induced stroke causes ischemia followed by reperfusion damage, mimicking the conditions of the most common type of stroke in human patients [1,10]. In this study, we induced a mild to moderate stroke in mice, ensuring that the animals remained ambulatory after the procedure, to characterize the effect of stroke beyond mechanical disuse effects on bone microstructure, dynamic bone formation, and tissue composition following four weeks of recovery. Since exercise initiated early during stroke recovery is associated with better motor control recovery and BMD maintenance [11–13], mice also performed daily treadmill locomotion. The goals of this study were 1) to characterize the effect of stroke without bedrest on bone parameters beyond BMD, and 2) to determine the effect of moderate daily exercise on stroke-induced bone changes.

## METHODS

### *in vivo* Assessments

#### Study Design

The protocol for this project was approved by the North Carolina State University Institutional Animal Care and Use Committee. Mice were housed by surgery group (4-5 per cage) on a 12-hour diurnal light cycle with access to chow and water *ad libitum*. Twenty-seven, 12-week-old, male, C57Bl/6J mice (The Jackson Laboratory, Bar Harbor, ME) received either a stroke (n = 15) or sham (n = 12) surgery (Fig. 1). Following surgery, mice were either given daily treadmill exercise (n = 8 stroke-exercise, n = 6 sham-exercise) or placed on a stationary treadmill for an equivalent time period (n = 7 stroke-sedentary, n = 6 sham-sedentary). For four days following surgery, mice were housed individually in cages with wetted food and hydrogel packs, and their health was monitored at least twice a day. Body mass was measured twice a day during the 4-day acute recovery period and weekly thereafter. On the fifth day, mice were returned to their original cages and group-housed. Stroke recovery and exercise therapy lasted for four weeks following surgery. For dynamic histomorphometry, fluorochrome labels were injected intraperitoneally using a 30 mg/kg (0.003 wt %) concentration at 10 days (alizarin complexone) and 3 days (calcein) prior to sacrifice. After four weeks, mice were euthanized with CO_2_ asphyxiation followed by cervical dislocation. Left and right femora and tibiae were collected, fixed in 10% neutral buffered formalin for 18 hours at 4°C (39.2°F), and then stored in 70% ethanol at 4°C (39.2°F) until analysis.

**Figure 1.**
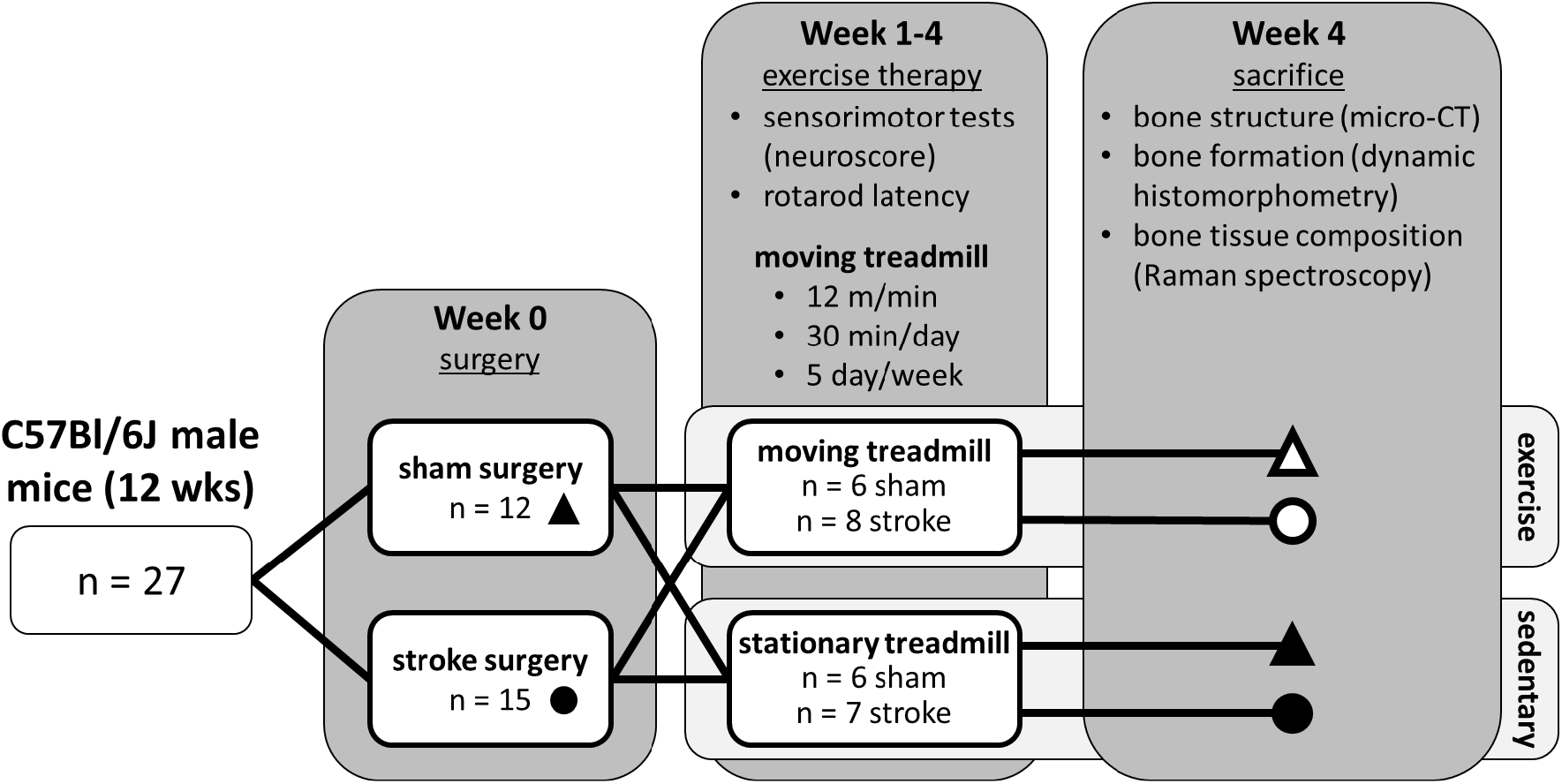
Experimental design: Twenty-seven C57Bl/6J mice received either a sham or stroke surgery at 12 weeks of age. Mice were further divided into either a moving treadmill exercise group or a stationary treadmill sedentary group. After four weeks of recovery, mice were sacrificed, and their femora and tibiae were collected for analysis.

#### Stroke Procedure

After mice were fasted 6-8 hours, ischemic stroke was induced in the right hemisphere using the well-established intraluminal middle cerebral artery occlusion (MCAo) model, in which a thin, silicone-coated 6-0 nylon filament is passed through the external carotid artery (ECA) and internal carotid artery (ICA) to the MCA origin, blocking blood flow to part of the ipsilateral cerebrum (mainly the cortex and basal ganglia) [10,14,15]. Mice were anesthetized with isoflurane (5% induction, then maintained around 2% throughout the surgery) in a 70:30 mixture of N_2_:0_2_ gas. Rectal temperature was maintained at 37°C (98.6°F) throughout the surgery with a heating pad (TCAT-2DF, Physitemp Instruments, LLC, Clifton, NJ). Cerebral blood flow (CBF) was monitored with laser Doppler flowmetry (moorVMS-LDF, Moor Instruments Ltd, Axminster, UK) using a monofilament probe (VP10M200ST, Moor Instruments Ltd). The probe was inserted through a small skin incision over the right temporal bone, just under the temporalis muscle, and affixed to the skull with a cyanoacrylate glue.

The MCAo and sham procedures were performed using aseptic technique. A midline skin incision was made along the neck, and the right common carotid artery (CCA), ECA, and ICA were exposed. For the stroke surgery, the CCA was temporarily ligated, the distal ECA and branches were ligated and cut, and the ICA was then temporarily ligated. At this point, the CCA ligation was temporarily loosened to record a pre-occlusion, baseline value of CBF. The CCA ligation was retightened, and the occluding filament (6-0 nylon filament with silicone-coated tip [1-3 mm (0.039-0.12 in) long, 0.22-0.23 mm (0.0087-0.0091 in) in diameter], Doccol Corporation, Redlands, CA) was passed through the stump of the ECA into the ICA until resistance was felt, indicating the occluding filament had reached the MCA origin. The filament was left in place for 30 minutes, and CBF was monitored to ensure an 80% CBF reduction throughout the occlusion period. Saline was added to the wound to ensure the tissue remained hydrated. After 30 minutes, the filament was removed, the ECA ligation was permanently tightened, and the CCA ligation was removed. For the sham surgery, after the neck incision was made and arteries exposed, saline was added to the neck incision, and CBF was monitored for 30 minutes without inserting the occluding filament. At the end of each surgery, the laser Doppler flowmetry probe was removed, an intra-incisional injection of bupivacaine (2 mg/kg, Marcaine, Hospira, Lake Forest, IL) was administered to the neck incision, and the neck and skull skin incisions were sutured closed. A 4% lidocaine cream and an antibiotic ointment were applied to the incision sites, and a subcutaneous injection of carprofen (Rimadyl, Zoetis, Parsippany, NJ) at 5 mg/kg (0.0005 wt %) was administered.

#### Sensorimotor Function

Ischemic stroke severity and functional recovery were assessed with a series of scored tests that were summed to form a neurological score, or *neuroscore*. Each test was scored from 0 to 2 or 0 to 4, depending on the test. The scores were determined empirically relative to the behavior seen with a sham surgery (score = 0) up to the behavior with severe stroke (score = 48, the maximum possible score). The scoring system was comprised of four main parts that were modified from similar systems used to assess sensorimotor function during stroke recovery in rats [16,17]: 1) mouse activity level, body symmetry, and gait were scored as the mice were allowed to ambulate freely in an open-top enclosure to assess gross impairments; 2) limb symmetry, circling, and limb placement were also scored during free ambulation to assess motor function; 3) climbing ability on a vertical metal screen and balance beam walking across a cylindrical rod were scored to determine complex motor function; and 4) sensory function was assessed by gently touching the left (paretic, affected) and right (non-paretic, unaffected) paws, trunk, vibrissae, and face of the mice with a cotton-tip applicator. Neuroscore examinations were performed at 24 hours, 48 hours, 3 days, 4 days, and then weekly following surgery.

Stroke recovery was also assessed using a rotarod, or rotating rod, test [15]. Mice were placed on a plastic rod that rotates at a velocity that accelerates from 4 to 40 rotations per minute over 5 minutes (ENV-576M, Med Associates Inc, St. Albans, VT). The test scores motor coordination by timing how long the mouse can walk on the rod without falling off or hugging the rod for three consecutive rotations without attempting to walk. Mice were acclimated to the rotarod test for two days immediately prior to surgery using a constant velocity of 20 rotations per minute. Mice performed accelerating rotarod tests at 48 hours, 4 days, and then weekly following surgery, on the same day as neuroscore testing. During each acclimation and rotarod test day, each mouse attempted the test three times, and the longest time was recorded, up to a maximum of 300 seconds. Mice were allowed 15 min of rest between each attempt.

#### Treadmill Exercise and Gait Pattern Analysis

Treadmill exercise was performed using a rodent treadmill (Exer 3/6, Columbus Instruments, Columbus, OH). Mice in the exercise groups were acclimated to the treadmill for two days prior to surgery at 6 m/min (6.56 yds/min) for 10 min. Exercise therapy began at four days after surgery, gradually increasing the protocol for the first two days. On day four, mice were exercised at 5 m/min (5.47 yds/min) for 10 min, followed by 12 m/min (13.12 yds/min) for 10 min. On day five, mice exercised at 5 m/min (5.47 yds/min) for 5 min, followed by 12 m/min (13.12 yds/min) for 20 min. Every weekday thereafter (5 days/wk) mice performed the full exercise routine of 12 m/min (13.12 yds/min) for 30 min. Sedentary group mice were placed on an immobile, replica treadmill for a matched time period to equalize the stress of animal handling across groups.

For the exercise groups only, interlimb coordination during treadmill gait was quantified weekly using video analysis. Altered gait in the affected hindlimb could alter the functional tissue strain experienced by bones during locomotion [18,19], which can affect bone formation [20,21]. Gait was captured over 60 s with high-speed video (240 frames/sec, HERO4, GoPro Inc, San Mateo, CA) as mice ran on the treadmill at 12 m/min (13.12 yds/min), and the videos were analyzed using Kinovea (version 0.8, Kinovea Open Source Project). Duty cycle between the hindlimbs and phase dispersion in relation to the left (affected) hindlimb were calculated based on the relative timing of gait pattern events [18,22,23]. Duty cycle, the ratio of stance times (paw strike to lift off) across a single gait cycle (paw strike to next paw strike), in the affected hindlimb was calculated relative to the duty cycle in the unaffected hindlimb. Phase dispersion, the relative timing of paw strikes between two limbs within the same gait cycle, was calculated between the affected hindlimb and each of the other limbs: unaffected hindlimb (contralateral), unaffected forelimb (diagonal), and affected forelimb (ipsilateral). All parameters were measured in three sets of five consecutive gait cycles for each weekly treadmill session, and the average parameters were calculated for each week. Video was captured at one day prior to surgery, five days after surgery, and weekly thereafter.

### Micro-Computed Tomography

Left and right femora were scanned in 70% ethanol with micro-computed tomography (μCT80, SCANCO Medical AG, Brüttisellen, Switzerland) using a 10-μm voxel size, 45 kV peak X-ray tube potential, 177 μA X-ray intensity, and 800-ms integration time. Volumes of interest (VOI) were analyzed in the distal metaphysis and mid-diaphysis. The metaphyseal VOI was defined as 10% of the total femur length, located proximal to the distal growth plate. The cancellous bone and cortical bone were contoured and analyzed separately in the metaphyseal VOI. Standard cancellous and cortical microstructural parameters were analyzed using the scanner’s software (SCANCO v.6.6) [24]. The diaphyseal VOI was defined as 15% of the total femur length, centered at the midpoint between the distal growth plate and middle of the third trochanter, and standard cortical bone parameters were analyzed.

### Dynamic Histomorphometry

Left and right tibiae were cut transversely at the tibiofibular junction under constant irrigation with water using a low-speed precision saw (IsoMet Low Speed Precision Cutter, Buehler, Lake Bluff, IL). The proximal bone segment was saved for Raman spectroscopy, and the distal segment was embedded in methylmethacrylate for dynamic histomorphometry [25]. In the distal segment, approximately 200-μm (0.0079-in) thick transverse sections were cut with the precision saw just distal to the tibiofibular junction, affixed to glass slides with cyanoacrylate glue, then sanded to 10-30 μm (0.00039-0.0012 in) thickness with increasing grit sandpaper. Two sections from each bone were imaged at 40X on a Zeiss LSM 880 laser scanning microscope with Airyscan (Carl Zeiss Microscopy, Thornwood, NY). Standard indices of dynamic bone formation were measured on two sections per limb using FIJI (built on ImageJ version 1.51n) and Photoshop (version CC 2018, Adobe Systems Inc., San Jose, CA) [26–28].

### Raman Spectroscopy

Cortical bone tissue composition was analyzed in the diaphysis of affected (left) tibiae using Raman spectroscopy. In the proximal bone segment described above, approximately 1-mm (0.39-in) thick sections were cut transversely using the Isomet Low Speed Precision Cutter just proximal to the tibiofibular junction. Sections were affixed to glass slides proximal side down with cyanoacrylate glue, then polished to create a smooth surface. Raman spectra were collected with a 785-nm laser at 50X magnification (Horiba XploRA PLUS, Jobin Yvon, Longjumeau, France). Each cortical section was scanned at the endosteal, mid-cortex, and periosteal edges on the anterior and posterior regions of the cross section (Fig. 2A). Each scan was comprised of a line of 6 points spaced 2 μm (7.87×10^−5^ in) apart, parallel to and 5-10 μm (1.97×10^−4^–3.94×10^−4^ in) in from the bone surface. Each point was a 30-second accumulation in the 800-1800 cm^−1^ range. Baseline correction was performed in LabSpec (v.6.5.1.24, HORIBA, Kyoto, Japan), and spectral analysis was performed with custom code in MATLAB^®^ (R2017, The MathWorks, Natick, MA).

**Figure 2.**
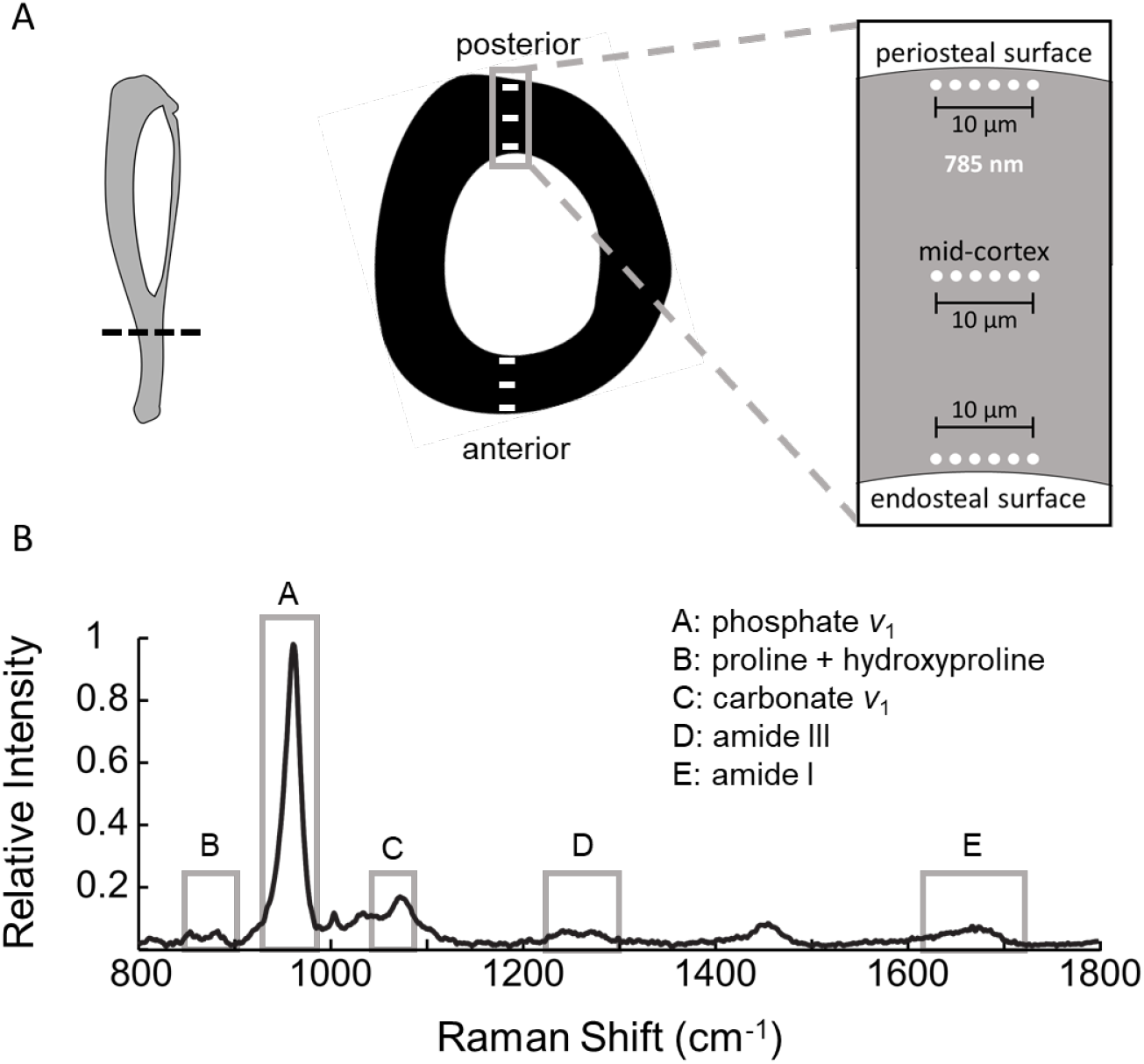
A) In cortical sections of the affected (left) tibial diaphysis, Raman spectra were collected at 6 equallyspaced regions, along a 10-μm long line on the anterior and posterior sides at three locations: endosteal edge, mid-cortex, and periosteal edge. B) Raman spectra were normalized to the phosphate phosphate *ν*_1_ band intensity, and band intensity ratios were calculated: phosphate *v*_1_/(proline+hydroxyproline), phosphate *v*_1_/carbonate *v*_1_ (carbonate substitution), phosphate *v*_1_/amide I, phosphate *v*_1_/amide III, and carbonate *v*_1_/amide I. Crystallinity was measured as the inverse of the full width at half-maximum (FWHM) of the phosphate *v*_1_ band.

Raman spectra were normalized relative to the maximum height (intensity) of the phosphate *v*_1_ band located at 930-980 cm^−1^, and maximum intensities relative to the phosphate *v*_1_ intensity were calculated for the following tissue constituents: summation of proline (830-863 cm^−1^) and hydroxyproline (wavenumber 864-899 cm^−1^), carbonate *v*_1_ (1055-1090 cm^−1^), amide I (1616-1720 cm^−1^), and amide III (1220-1300 cm^−1^) [29,30] (Fig. 2B). Several standard band intensity ratios were calculated [29,31]. Mineral-to-matrix ratios were calculated using phosphate relative to several matrix bands: amide I, amide III, or summed proline and hydroxyproline. Carbonate substitution was calculated as the band intensity ratio of carbonate *v*_1_ relative to phosphate *v*_1_, and the carbonate-to-matrix ratio was calculated as carbonate *v*_1_ relative to amide I. Mineral maturity (crystallinity) was calculated as the inverse of the full width at half maximum (FWHM) of a single-order Gaussian curve fit to the phosphate *v*_1_ band.

### Statistical Analysis

Statistical models were analyzed in SAS (SAS University Edition v. 9.4, SAS Institute Inc., Cary, NC) to determine the following: 1) effects of stroke and exercise therapy on neuroscore, rotarod time, and body mass at each timepoint; 2) effect of stroke on gait parameters at each timepoint; 3) effects of stroke and exercise on femoral microstructure, tibial dynamic bone formation, and tibial composition; and 4) whether stroke differentially affects femoral microstructure and tibial bone formation between affected and unaffected limbs.

For analysis #1, stroke recovery parameters (neuroscore, rotarod, body mass) were compared between surgery and activity groups at each timepoint using a repeated measures factorial model (procedure MIXED) with interaction between all terms. Surgery group (sham or stroke) and activity group (sedentary or exercise) were modeled as fixed factors, while timepoint was modeled as a repeated measure within each mouse. The residual variance was modeled assuming compound symmetry covariance. Predicted least-squares means with Tukey-Kramer adjustment for multiple comparisons were used to analyze effect differences between surgery and activity groups, with interaction, within each timepoint (i.e., sham-sedentary vs. stroke-sedentary at Week 3). Analysis #2 used a similar model, except activity group was not included, since gait was only analyzed in the exercise groups (i.e., sham vs. stroke at Week 3).

Analyses #3 and 4 also used a repeated measures factorial model (procedure MIXED). Parameters of femoral microstructure and tibial histomorphometry were analyzed between activity and surgery groups across limb (repeated measure). Tibial composition was only analyzed in the affected tibiae, but scan region (anterior and posterior) was modeled as the repeated measure. Tibial composition in the endosteal, mid-cortex, and periosteal regions were each run as separate models and were not compared. All models used compound symmetry to calculate residual variance. For analysis #3 – the effect of activity, stroke, and their interaction on femoral and tibial bone parameters – predicted least-squares means were used to perform pairwise differences (i.e., sham-sedentary vs. stroke-sedentary) with Tukey-Kramer adjustments for multiple comparisons. For analysis #4, least-squares means were used to analyze differences between limb across surgery group (i.e., affected vs. unaffected limb within stroke group).

A significance level of 0.05 was used for all analyses. Data are presented as mean ± standard deviation unless otherwise noted. Data from methods that analyzed affected and unaffected limb bones (i.e., femoral microstructure and tibial histomorphometry) are presented as the mean for both limbs per mouse. Results from Raman spectroscopy are presented as the mean across anterior and posterior quadrants per bone.

## RESULTS

### *in vivo* Assessments

Following stroke, all neuroscores were above 0 (sham group score) at all post-stroke timepoints, confirming that the MCAo procedure caused neurological impairment (Fig. 3A). Based on these neuroscores, the severity of the induced strokes was mild to moderate, and the magnitude of impairment varied between individual mice and tended to improve over time (decreasing neuroscore). Exercise did not affect neuroscore at any timepoint (p = 0.66), suggesting the treadmill exercise protocol used did not affect stroke recovery. The rotarod test was not a useful metric for assessing motor impairments post-stroke, as it did not capture the same differences as the neuroscore, with no significant effects of stroke (p = 0.88) or exercise (p = 0.37) observed at any timepoint (Fig. 3B). Before surgery, sham and stroke groups had similar body mass (p = 0.38), but the stroke group experienced significant mass losses after ischemic stroke and remained smaller than the sham group at all timepoints post-surgery (Fig. 3C).

**Figure 3.**
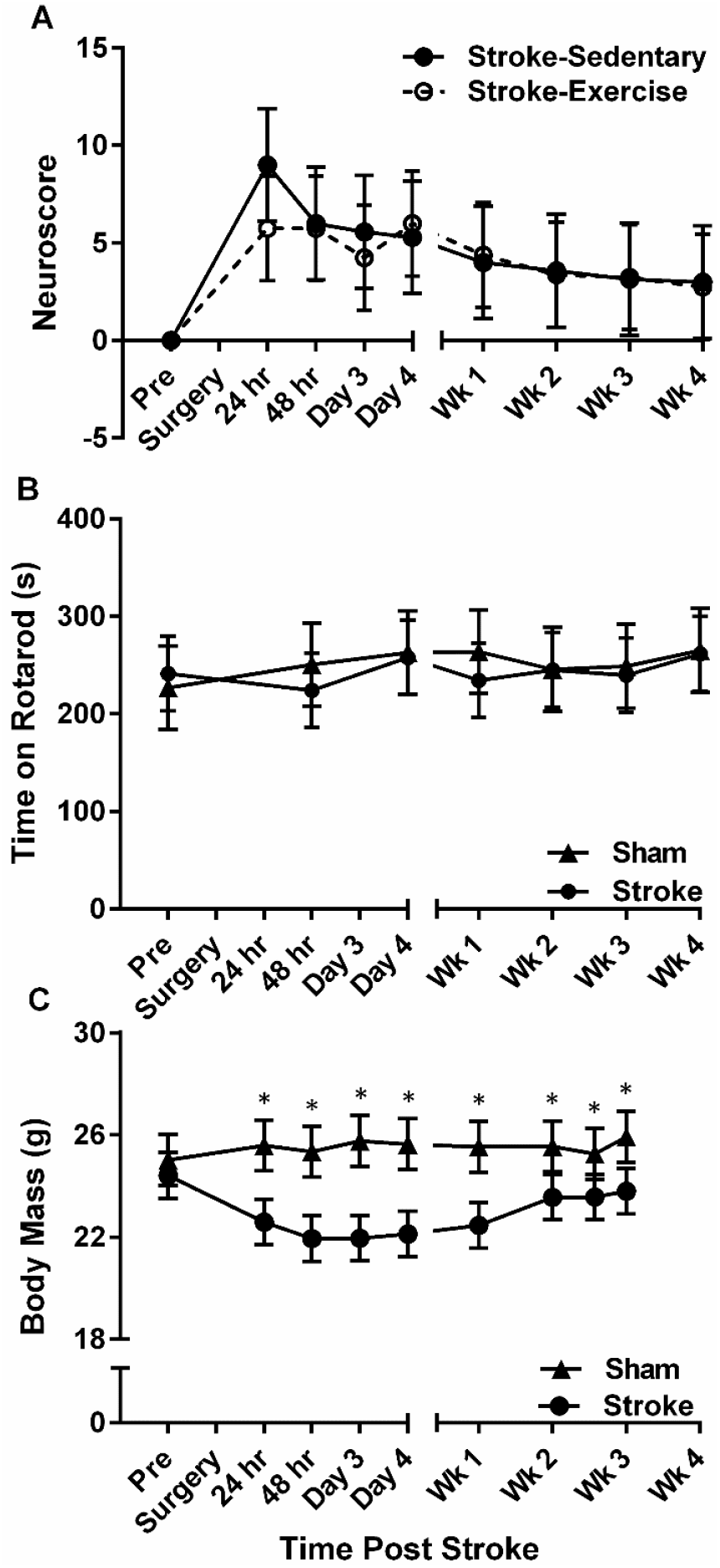
Stroke severity and recovery were assessed with A) scored tests of sensorimotor function (*neuroscore*), B) length of time spent on an accelerating rod, and C) body mass. Data presented as estimated least-squares means ± 95% confidence intervals. *p < 0.05 stroke vs. sham within timepoint.

### Gait Pattern Analysis

Limb coordination patterns were quantified in the exercise groups only to avoid confounding the sedentary groups with treadmill activity. The stroke-exercise group was able to perform the daily treadmill regimen beginning four days after stroke. Gait parameters involving the affected (left) hindlimb were mostly similar between stroke and sham groups at every timepoint (Fig. 4). Diagonal phase dispersion was nearly lower in stroke compared to sham groups in Week 1 (16%, p = 0.067) and Week 2 (3.4%, p = 0.085) (Fig. 4B), but both contralateral (Fig. 4C) and ipsilateral (Fig. 4D) phase dispersions did not differ significantly between stroke and sham groups at any timepoint. The shift in diagonal phase dispersion represents the affected hindlimb paw strike occurring earlier relative to the right forelimb paw lift, which could be a slight limp in the affected hindlimb. However, the duty cycle ratio between the hindlimbs was unaffected by stroke at any timepoint (Fig. 4A), suggesting no differences in hindlimb favorability between sham and stroke groups. Together, these data suggest that limb loading, and thus functional bone tissue strain, during exercise was similar for both groups. One outlier was removed from the sham-exercise group in Week 1, because the limb coordination patterns were irregular, as the mouse was sprinting and pausing excessively and not running at the constant velocity of the treadmill. Several timepoints had videos that could not be analyzed due to poor lighting or camera placement. Except for Week 2, where only two videos were analyzed for sham-exercise, at least four videos were analyzed per group per timepoint.

**Figure 4.**
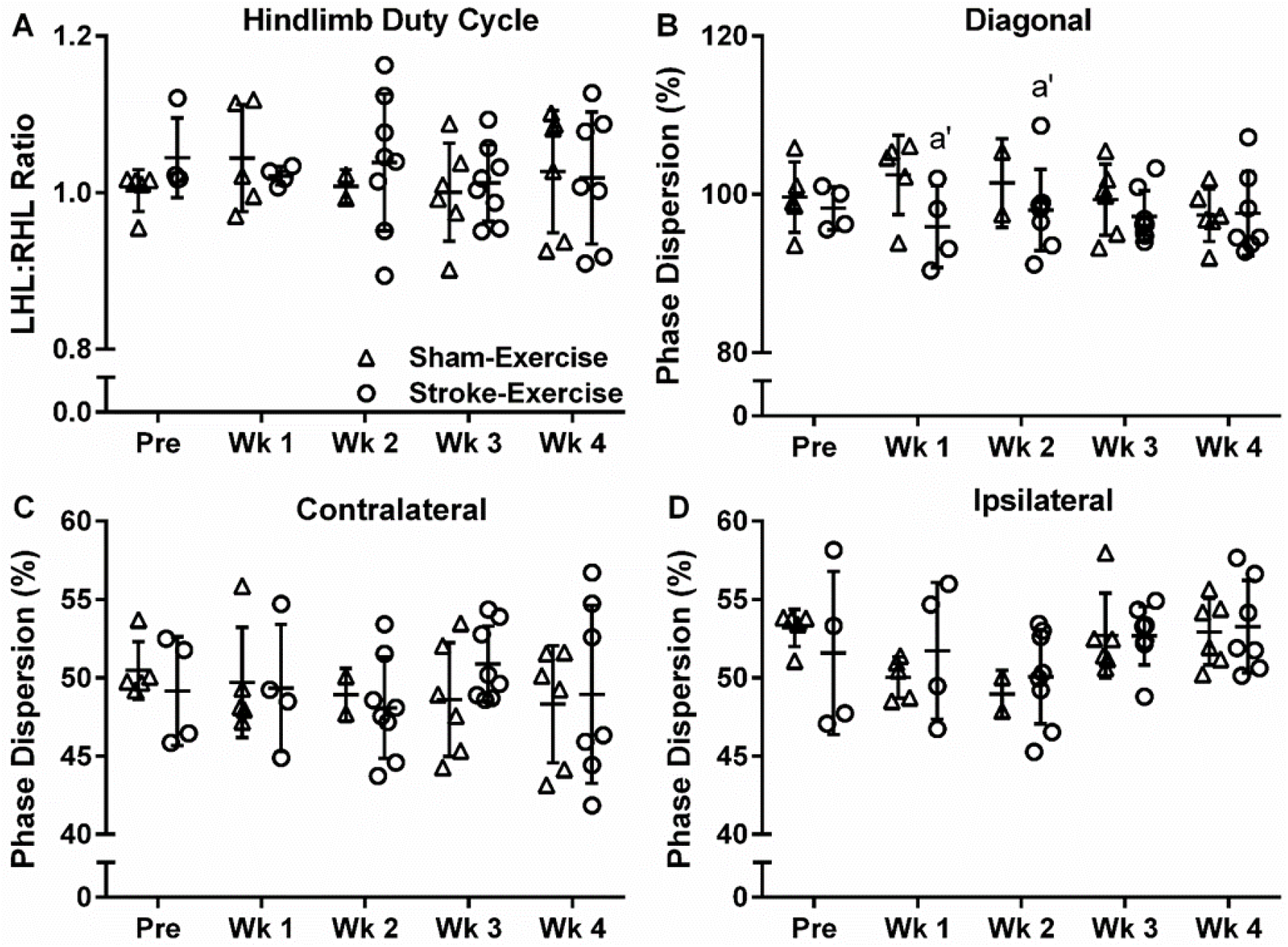
Limb coordination parameters involving the affected left hindlimb during treadmill locomotion were all unaffected by stroke: A) duty cycle ratio between affected left hindlimb (LHL) and unaffected right hindlimb (RHL), B) diagonal phase dispersion, C) contralateral phase dispersion, and D) ipsilateral phase dispersion. a’: p < 0.1 sham-exercise vs. stroke-exercise within timepoint

### Micro-Computed Tomography

In all VOIs, stroke did not affect femoral microstructure (stroke-sedentary vs. sham-sedentary), but exercise improved femoral microstructure in the sham group (sham-exercise vs. sham-sedentary) but not the stroke group (stroke-exercise vs. stroke-sedentary). In the metaphysis, cortical area (Ct.Ar) was nearly greater in sham-exercise than sham-sedentary (11%, p = 0.066) and was 17% greater than stroke-exercise (p = 0.0032) but was similar between stroke-exercise and stroke-sedentary (p = 0.45) (Fig. 5A). Similarly, total area (Tt.Ar) was greater in sham-exercise than stroke-exercise (12%, 3.40 ± 0.19 mm^2^ vs. 3.03 ± 0.28 mm^2^, p = 0.046) but was similar between sham-exercise and sham-sedentary (3.12 ± 0.06 mm^2^, p = 0.24), and between stroke-exercise and stroke-sedentary (3.16 ± 0.14 mm^2^, p = 0.69). Ct.Ar and Tt.Ar were similar between stroke-sedentary and sham-sedentary (p = 1.00 and p = 0.99, respectively). Bone mineral density (BMD), tissue mineral density (TMD), cortical thickness (Ct.Th), and cortical area fraction (Ct.Ar/Tt.Ar) were not affected by exercise or stroke. In the diaphysis, Ct.Ar was 13-18% greater in sham-exercise compared to all other groups (p < 0.0001 to p = 0.0039) but was similar between stroke-exercise and stroke-sedentary (p = 0.64) and between stroke-sedentary and sham-sedentary (p = 0.97) (Fig. 5D). Tt.Ar was nearly greater in sham-exercise than sham-sedentary (13%, 2.28 ± 0.21 vs. 2.01 ± 0.06 mm^2^, p = 0.085) and was 20% greater than stroke-exercise (vs. 1.90 ± 0.14 mm^2^, p = 0.0046) but was similar between stroke-exercise and stroke-sedentary (vs. 2.03 ± 0.12 mm^2^, p = 0.44) and between stroke-sedentary and sham-sedentary (p = 1.00). BMD, TMD, Ct.Th (Fig. 5E), and Ct.Ar/Tt.Ar (Fig. 5F) were not affected by exercise or stroke.

**Figure 5.**
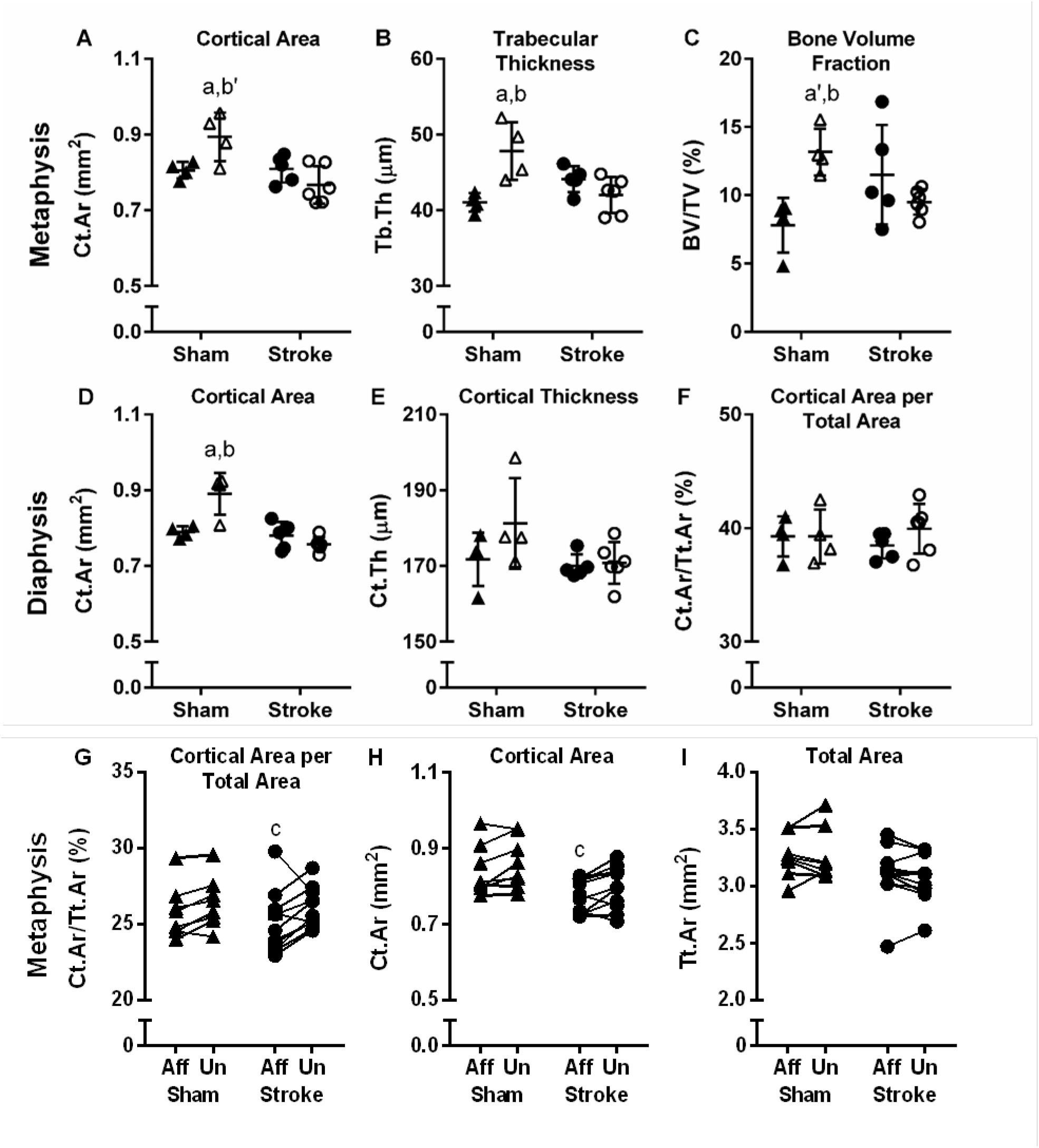
Femoral microstructure was assessed for group differences in the distal metaphysis (A-C) and mid-diaphysis (D-F) and for affected-unaffected limb differences in the distal metaphysis (G-I). In the metaphysis, exercise in sham, but not stroke, caused increased A) cortical area, B) trabecular thickness, and C) bone volume fraction. In the diaphysis, D) Ct.Ar also increased with sham-exercise relative to sham-sedentary and stroke-exercise; E) cortical thickness and F) cortical area per total area were unaffected by stroke or exercise. In the metaphysis, affected-to-unaffected limb differences were observed in stroke but not sham groups for G) Ct.Ar/Tt.Ar and H) Ct.Ar but not I) Tt.Ar. a: p < 0.05 sham-exercise vs. stroke exercise. a’: p < 0.1 sham-exercise vs. stroke exercise. b: p < 0.05 sham-exercise vs. sham-sedentary. b’: p < 0.1 sham-exercise vs. sham-sedentary. c: p < 0.05 vs. unaffected side within surgery group.

Similar trends of no stroke effect but inhibited exercise-induced microstructural gains in the stroke group were observed in the cancellous bone of the distal metaphysis. Trabecular thickness (Tb.Th) was greater in sham-exercise than both sham-sedentary (17%, p = 0.0060) and stroke-exercise (14%, p = 0.010), but stroke-exercise was similar to stroke-sedentary (p = 0.51) (Fig. 5B). Bone volume fraction (BV/TV) was 69% greater in sham-exercise than sham-sedentary (p = 0.022) and nearly greater than stroke-exercise (38%, p = 0.10), but stroke-exercise was similar to stroke-sedentary (p = 0.49) (Fig. 5C). Degree of anisotropy (DA) was nearly greater in sham-exercise than sham-sedentary (11%, 1.49 ± 0.05 vs. 1.34 ± 0.06, p = 0.089) and was 13% greater than stroke-exercise (vs. 1.33 ± 0.08, p = 0.031), but stroke-exercise was similar to stroke-sedentary (vs. 1.43 ± 0.11, p = 0.24). Stroke-sedentary was similar to sham-sedentary for Tb.Th (p = 0.27), BV/TV (p = 0.12), and DA (p = 0.47). BMD, TMD, trabecular number (Tb.N) and separation (Tb.Sp), and connectivity density (Conn.D) were not affected by exercise or stroke. Although stroke did not cause detrimental changes to cancellous microstructure, exercise improved several microstructural parameters in the sham group only, suggesting stroke may inhibit bone anabolism even in ambulatory mice.

Stroke caused affected-unaffected side differences within individual mice (paired comparisons) in some microstructural parameters. In the cortical bone surrounding the distal metaphysis Ct.Ar/Tt.Ar was 1% smaller (p = 0.019, Fig. 5G) and Ct.Ar was 2% smaller (p = 0.032, Fig. 5H) in the affected side compared to the unaffected side for the stroke group, but Tt.Ar was similar between stroke affected and unaffected sides (p = 0.27, Fig. 5I). Cortical BMD was 4% greater in the affected side compared to the unaffected side for the stroke group (633 ± 23 vs. 611 ± 25 mg/cm^3^, p = 0.0018) but similar between the affected and unaffected side for the sham group (634 ± 17 vs. 632 ± 16 mg/cm^3^, p = 0.74). Metaphyseal trabecular parameters and diaphyseal cortical parameters were not significantly different between affected and unaffected sides for the stroke group. Similarly, no affected-unaffected side differences were found in the sham group, except trabecular separation, which was 3% bigger in the affected than unaffected side (238 ± 34 vs. 232 ± 27 μm, p = 0.035).

### Dynamic Histomorphometry

Stroke had no effect on dynamic indices of bone formation – mineralizing surface per bone surface (MS/BS), mineral apposition rate (MAR), and bone formation rate (BFR) – on either the periosteal or endosteal surface in the tibial midshaft (Fig. 6). On the endosteal surface, exercise decreased MS/BS by 25% relative to sedentary (p = 0.0078, Fig 6A), had no effect on MAR (p = 0.58, Fig. 6B), and nearly decreased BFR (14%, p = 0.090, Fig. 6C). Although MS/BS did not differ significantly between affected and unaffected tibiae for either stroke or sham groups (Fig. 6D), MAR was 40% lower in the affected than the unaffected tibia for the stroke group (p < 0.0001) and 30% lower in the sham group (p = 0.0045) (Fig. 6E). Similarly, BFR was 44% lower in the affected tibia relative to the unaffected tibia for the stroke group (p = 0.0005) and nearly lower in the sham group (36%, p = 0.10) (Fig. 6F). On the periosteal surface, exercise had no effect on MS/BS, MAR, or BFR. The sample size for the sham-exercise group was small (n = 2) across all of these analyses, because one cage of mice in this group missed the alizarin complexone injection. Therefore, only main effects (exercise, activity) were examined for these parameters, and more samples are needed to confirm the exercise effect observed in the sham-exercise group.

**Figure 6.**
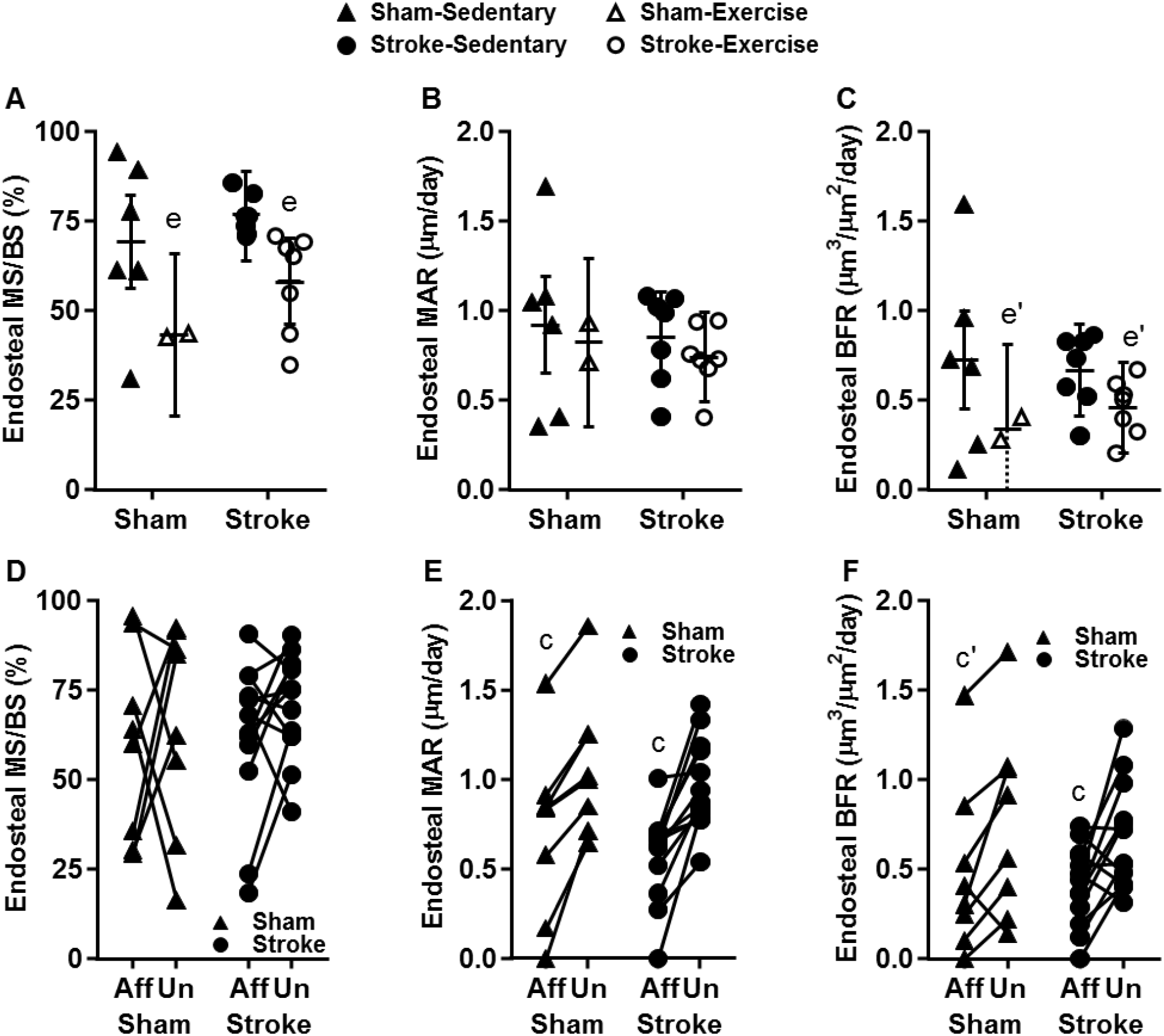
Dynamic cortical bone formation was assessed in the tibial diaphysis with dynamic histomorphometry. No differences were found on the periosteal surface. On the endosteal surface, exercise decreased (relative to sedentary) A) mineralizing surface per bone surface (MS/BS) but not B) mineral apposition rate (MAR) or C) bone formation rate (BFR). For affected (Aff) vs. unaffected (Un) limb differences, neither sham nor stroke group showed D) differences for MS/BS, but E) both sham and stroke had decreased MAR in the affected limbs, and F) stroke decreased BFR in the affected limb. (A-C) Data presented as estimated least-squares mean ± 95% confidence interval. c: p < 0.05 vs. unaffected side within surgery group. d: p < 0.05 exercise vs. sedentary (main effect).

### Raman Spectroscopy

Stroke induced minor changes in bone tissue composition in the diaphyseal bone of affected tibiae, while exercise induced more substantial changes, particularly in the stroke group (Fig. 7). Exercise increased phosphate:amide III mineral-to-matrix ratio in stroke-exercise relative to stroke-sedentary by 27% near the endosteal surface (p = 0.019) and by 25% in the mid-cortex (p = 0.0070) but not in sham-exercise relative to sham-sedentary (p = 1.00 endosteal, p = 0.58 mid-cortex) (Fig. 7A). Stroke nearly decreased phosphate:amide III mineral-to-matrix in stroke-sedentary relative to sham-sedentary near the endosteal surface (18%, p = 0.10) but not in the midcortex (p = 0.91). Exercise also increased the endosteal phosphate:amide I mineral-to-matrix ratio (23%, p = 0.079, Fig. 7C) and the endosteal carbonate-to-matrix ratio (20%, p = 0.11, Fig. 7D) in stroke-exercise relative to stroke-sedentary, but not in sham-exercise relative to sham-sedentary (p = 0.96 and p = 0.80, respectively). Exercise had no effect on the phosphate:(proline+hydroxyproline) mineral-to-matrix ratio (Fig. 7B).

**Figure 7.**
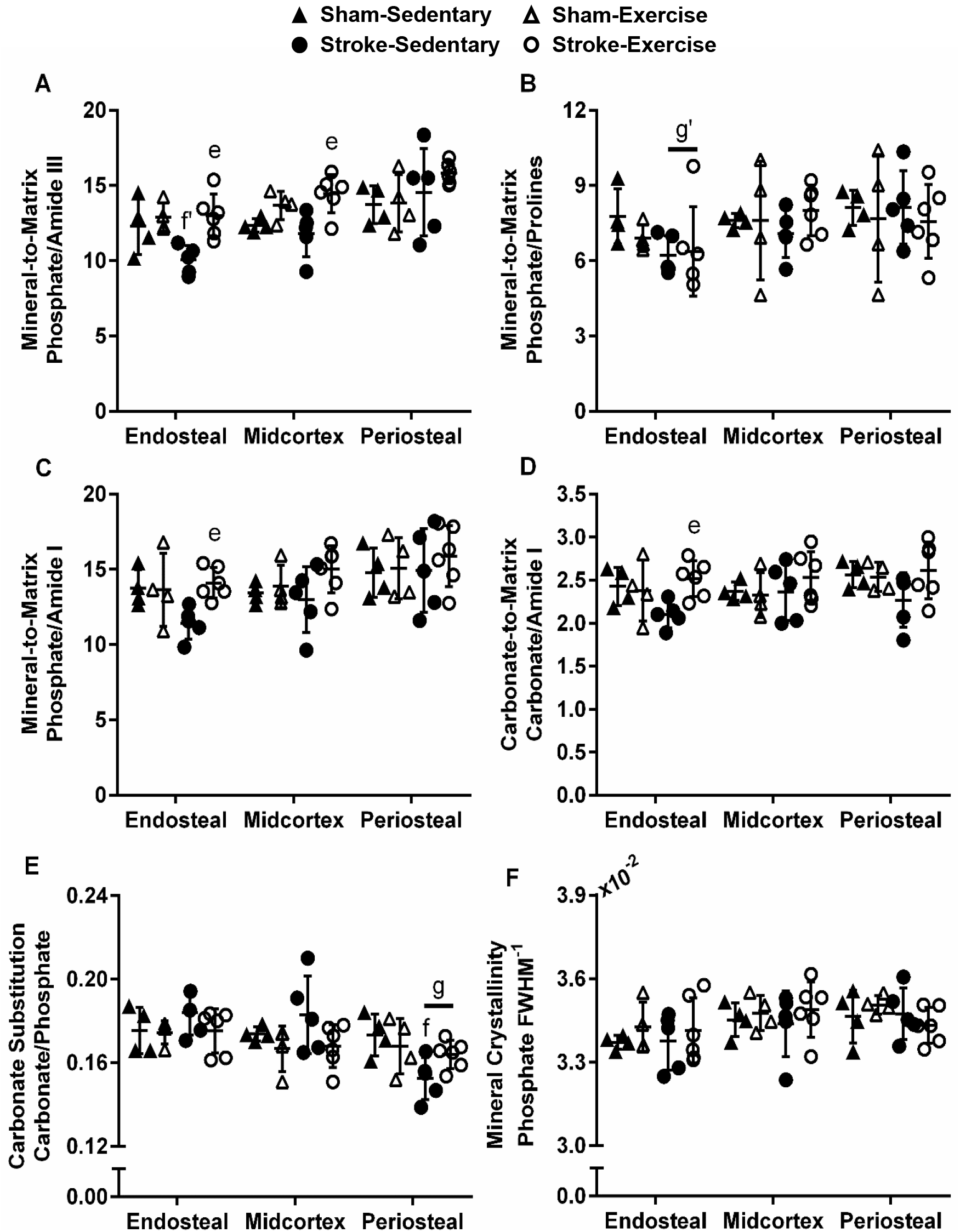
Bone tissue composition assessed in the endosteal, mid-cortex, and periosteal regions of the affected tibial diaphysis revealed significant changes with exercise relative to sedentary and minor changes with stroke relative to sham: A) phosphate *v*_1_/amide III, B) phosphate *v*_1_/(proline+hydroxyproline), C) phosphate *v*_1_/amide I, D) carbonate *v*_1_/amide I, E) carbonate substitution (carbonate *v*_1_/phosphate *v*_1_), and F) mineral crystallinity (phosphate *v*_1_ full width at half maximum, FWHM). e: p < 0.05 stroke-exercise vs. stroke-sedentary. f: p < 0.05 stroke-sedentary vs. sham-sedentary. g: p <0.05 stroke vs sham (main effect).

Stroke decreased carbonate substitution near the periosteal surface by 8% relative to sham (main effect p = 0.019, Fig. 7E) and by 12% in stroke-sedentary relative to sham-sedentary (p = 0.029). Stroke also nearly decreased phosphate:(proline+hydroxyproline) mineral-to-matrix ratio near the endosteal surface (14%, p = 0.090, Fig. 7B). For carbonate substitution, carbonate-to-matrix ratio, and the phosphate:amide I and phosphate:amide III mineral-to-matrix ratios, the stroke-sedentary group exhibited the most differences relative to the other groups, while stroke-exercise was similar to sham-exercise, suggesting exercise may mitigate the effects of stroke on bone composition. Mineral maturity (crystallinity) was unaffected by stroke or exercise (Fig. 6F).

## DISCUSSION

Ischemic stroke in mice inhibited the anabolic effects of moderate treadmill exercise on femoral bone microstructure. The prevention of exercise-induced adaptations following stroke occurred in the absence of altered gait patterns, and thus despite a normal (sham-level) amount of limb loading. These results directly challenge the current paradigm that stroke effects on bone result primarily from mechanical unloading with bedrest and rather suggest that stroke has direct effects on bone that are independent of disuse effects. In particular, our results show that cortical and trabecular microstructure in the stroke-exercise group was similar to that in both the stroke-sedentary and sham-sedentary groups, while the sham-exercise group experienced metaphyseal and diaphyseal cortical bone enlargement and trabecular thickening that resulted in a more anisotropic orientation on average. Despite the urgent clinical need to understand the high fracture risk in human stroke patients, to our knowledge this study is only the second one to examine changes to bone tissue in an induced stroke model [32], and the first study in mice. This work reveals that the middle cerebral artery occlusion model of ischemic stroke in mice impacts bone microstructure, bone formation rate, and tissue composition without bedrest; further work with this model can be used to examine mechanisms underlying microstructural changes and inhibited exercise adaptation, which could inform future clinical studies.

The effects of stroke on bone were primarily systemic, with similar changes in both affected and unaffected limbs. Stroke did not change gait locomotion patterns in the affected hindlimb compared to sham and, therefore, the stroke and sham groups likely experienced similar functional bone tissue strain, which is known to affect bone remodeling [18–20,33]. Although diagonal phase dispersion was nearly lowered during the first two weeks post-stroke in stroke-exercise relative to sham-exercise, the amount of time spent in stance (duty cycle ratio) was similar between affected and unaffected hindlimbs. Therefore, the functional tissue strain was likely similar as well. Gait patterns were not analyzed in the sedentary groups, since high-speed video had to be captured during treadmill exercise, and even a small amount of exercise could confound the sedentary results. The acute recovery period appears to be a critical window during which physical activity could prevent skeletal health decline in human stroke patients [34]. Based on a pilot study, we found that the earliest that mice could perform the full treadmill regimen was beginning at four days after stroke. For earlier intervention, alternative physical therapies or exercise-mimicking pharmacological therapies would need to be explored. Future studies could also implement exercise training prior to MCAo, which has been shown to improve functional recovery following stroke [35].

Despite mostly systemic effects, stroke did induce differential changes in the affected limb (relative to the unaffected limb) in cortical bone microstructure and indices of bone formation. Similarly, another MCAo study in 11-week-old male rats that also found no changes in femoral microstructure, reported affected-unaffected limb differences in the stroke group [32]. They found that, in the cortical bone surrounding the distal femoral metaphysis, stroke increased total area (reported as total volume) by 2% and BMD by 5% in the affected relative to the unaffected side, while we found a 4% reduction in total area and a similar 3% increase in BMD. The difference in our total area could be due to stroke interacting with skeletal growth, since 11-week-old rats are at an earlier point in skeletal development than 12-week-old mice. Skeletal growth in C57Bl/6J mice slows at 12 weeks and reaches skeletal maturity at 16 weeks, while growth in rats slows around 15 weeks and reaches skeletal maturity at 20 weeks [36–38]. Since animals were ambulatory following stroke in both experiments, these results suggest that stroke, without bedrest, specifically affects the metaphyseal cortical bone envelope. In the tibial diaphysis, stroke differentially decreased mineral apposition rate and bone formation rate by nearly half relative to the unaffected side. However, a similar unaffected-affected side difference in mineral apposition rate was present in sham, as well. These results are consistent with the MCAo study in rats, which found decreased serum concentration of P1NP, a marker of bone formation, four weeks after stroke [32]. Although this outcome was observed in a different bone and site, stroke-induced decreased bone formation rate may contribute to the affected-unaffected limb differences in cortical area fraction observed in the femoral metaphysis.

Although exercise did not affect femoral microstructure in the stroke group, it significantly altered tissue composition, an important contributor to bone strength [29,39]. Along the endosteal surface of the tibial diaphysis of the affected limb, the stroke-exercise group had higher phosphate:amide III and phosphate:amide I mineral-to-matrix ratios and carbonate-to-matrix ratio compared to stroke-sedentary. Since minimal affected-to-unaffected limb differences were observed in other metrics, only affected side tibiae were analyzed. Increased mineral-to-matrix and carbonate-to-matrix ratios indicate stiffer, but potentially more brittle, bone that has not been recently remodeled [29,31]. However, increased mineralization may also be a mechanism by which bone adapts to mechanical load without changing tibial morphology [20,31,40]. A study with the same daily treadmill regimen in male C57Bl/6J mice, initiated at 16 weeks-of-age for three weeks, found no changes to tibial morphology but a 15% higher phosphate:(proline+hydroxyproline) mineral-to-matrix ratio and approximately 10% higher ultimate tensile strain in the tibia with exercise relative to sedentary [40]. Similarly, a study in young and old rats found that the phosphate:amide I mineral-to-matrix ratio was positively correlated with stiffness but negatively correlated with bending modulus in the femur [31]. Taken together with the dynamic histomorphometry results (lower endosteal mineralizing surface per bone surface with exercise, decreased mineral apposition rate in the stroke-affected limb), the increased tissue mineralization observed in the endosteal surface of stroke-exercise relative to sham-exercise may be compensatory to maintain functional bone tissue strain.

Exercise may also impact bone formation, evidenced here by a tendency for reduced mineralizing surface per endosteal bone surface. Treadmill exercise has been shown in other studies to stimulate bone adaptation in mice, although the effect is dependent on the bone envelope, age, sex, and mouse strain – tibiae from younger male mice tend have a greater anabolic response to treadmill exercise than tibiae from older mice past skeletal maturity [33,40–42]. Our results showed no clear anabolic effect with exercise (e.g., no change in bone formation rate and reduced mineralizing surface per surface relative to sham endosteally), but more samples from the sham-exercise group are needed to confirm this observation. However, two recent studies using a similar treadmill regimen in male C57Bl/6J of similar age also found that exercise had no effect on morphology of the tibial diaphysis or on endosteal bone formation rate, but did increase periosteal bone formation rate in the tibial diaphysis and bone volume fraction in the tibial metaphysis relative to sedentary [33,41]. Together, these studies closely mirror our observed exercise-induced increases to trabecular bone volume fraction in the distal femur and lack of change in dynamic indices of bone formation in the tibia, at least for the endosteal surface.

This study revealed that stroke, without bedrest, prevented exercise-induced gains in microstructure, caused affected-unaffected side differences in microstructure and bone formation rate, and affected bone tissue composition. The MCAo mouse model had several advantages over human clinical studies: bone microstructure, dynamic bone formation, and bone composition are difficult to assess in human patients; mice remain ambulatory following stroke, enabling study of the effects of stroke beyond disuse from bedrest; comorbidities preceding ischemic stroke in human patients are not present in mice; and stroke conditions (e.g., location, severity) are tightly controlled. The mouse MCAo model in this study did not cause BMD loss, as seen in human stroke patients, but did cause detriments to microstructure in the affected limb. In the future, the study design could be adapted to try to recapitulate BMD loss seen in human stroke by increasing stroke severity (i.e., occluding the artery for a longer period), increasing the age of the mice, incorporating hindlimb suspension (to mimic bedrest), or extending the endpoint for a longer recovery period. Stroke causes chronic systemic inflammation [43,44], declining vascular health [45], and damage to the central nervous system, each of which are known to affect bone remodeling and maintenance [46,47]. The MCAo model is an ideal system to explore these interactions and their effects on skeletal health; lifestyle disease models and genetic modifications can be readily generated in mice, the conditions of the stroke are reproducible and easily manipulated, and different interventions can be implemented to mimic rehabilitation strategies. Improving our understanding of the factors contributing to the reduced skeletal adaptive response following stroke will be essential for developing better strategies to mitigate bone fragility in these patients.

## ACKNOWLEDGMENTS

We thank Dr. Ted Bateman and Eric Livingston for micro-CT support; Dr. Roberto Garcia, Dr. Chuanzhen Elaine Zhou, and Dr. Fred Stevie for Raman spectroscopy and sample preparation support; Dr. Eva Johannes for confocal microscopy support; John Biondi and Sophia Tushak for help with dynamic histomorphometry; and Dr. Consuelo Arellano for statistical consulting.

This work was performed in part at the Analytical Instrumentation Facility (AIF) at North Carolina State University, which is supported by the State of North Carolina and the National Science Foundation (award number ECCS-1542015). The AIF is a member of the North Carolina Research Triangle Nanotechnology Network (RTNN), a site in the National Nanotechnology Coordinated Infrastructure (NNCI). The authors acknowledge the use of the Cellular and Molecular Imaging Facility (CMIF) at North Carolina State University, which is supported by the State of North Carolina and the National Science Foundation.

## FUNDING

Research reported in this publication was supported by the Eunice Kennedy Shriver National Institute of Child Health and Human Development (NICHD) of the National Institutes of Health (NIH) under award number K12HD073945 and by the American Heart Association (AHA) under award number 7GRNT33710007. The content is solely the responsibility of the authors and does not necessarily represent the official views of the National Institutes of Health.

## REFERENCES

[1] Benjamin, E. J., Muntner, P., Alonso, A., Bittencourt, M. S., Callaway, C. W., Carson, A. P., Chamberlain, A. M., Chang, A. R., Cheng, S., Das, S. R., Delling, F. N., Djousse, L., Elkind, M. S. V., Ferguson, J. F., Fornage, M., Jordan, L. C., Khan, S. S., Kissela, B. M., Knutson, K. L., Kwan, T. W., Lackland, D. T., Lewis, T. T., Lichtman, J. H., Longenecker, C. T., Loop, M. S., Lutsey, P. L., Martin, S. S., Matsushita, K., Moran, A. E., Mussolino, M. E., O’Flaherty, M., Pandey, A., Perak, A. M., Rosamond, W. D., Roth, G. A., Sampson, U. K. A., Satou, G. M., Schroeder, E. B., Shah, S. H., Spartano, N. L., Stokes, A., Tirschwell, D. L., Tsao, C. W., Turakhia, M. P., VanWagner, L. B., Wilkins, J. T., Wong, S. S., Virani, S. S., and American Heart Association Council on Epidemiology and Prevention Statistics Committee and Stroke Statistics Subcommittee, 2019, “Heart Disease and Stroke Statistics-2019 Update: A Report From the American Heart Association,” Circulation, 139(10), pp. e56–e528. DOI: 10.1161/CIR.0000000000000659.

[2] Jones, G., Nguyen, T., Sambrook, P., Kelly, P. J., and Eisman, J. A., 1994, “Progressive Loss of Bone in the Femoral Neck in Elderly People: Longitudinal Findings from the Dubbo Osteoporosis Epidemiology Study.,” BMJ, 309(6956), pp. 691–695.

[3] Jørgensen, L., Jacobsen, B. K., Wilsgaard, T., and Magnus, J. H., 2000, “Walking after Stroke: Does It Matter? Changes in Bone Mineral Density Within the First 12 Months after Stroke. A Longitudinal Study,” Osteoporos Int, 11(5), pp. 381–387. DOI: 10.1007/s001980070103.

[4] Ramnemark, A., Nyberg, L., Lorentzon, R., Englund, U., and Gustafson, Y., 1999, “Progressive Hemiosteoporosis on the Paretic Side and Increased Bone Mineral Density in the Nonparetic Arm the First Year after Severe Stroke,” Osteoporos Int, 9(3), pp. 269–275. DOI: 10.1007/s001980050147.

[5] Borschmann, K., Pang, M. Y. C., Iuliano, S., Churilov, L., Brodtmann, A., Ekinci, E. I., and Bernhardt, J., 2015, “Changes to Volumetric Bone Mineral Density and Bone Strength after Stroke: A Prospective Study,” Int J Stroke, 10(3), pp. 396–399. DOI: 10.1111/ijs.12228.

[6] Beaupré, G. S., and Lew, H. L., 2006, “Bone-Density Changes After Stroke,” American Journal of Physical Medicine & Rehabilitation, 85(5), pp. 464–472. DOI: 10.1097/01.phm.0000214275.69286.7a.

[7] Jørgensen, L., and Jacobsen, B. K., 2001, “Functional Status of the Paretic Arm Affects the Loss of Bone Mineral in the Proximal Humerus after Stroke: A 1-Year Prospective Study,” Calcif. Tissue Int., 68(1), pp. 11–15. DOI: 10.1007/s002230001165.

[8] Ramnemark, A., Nyberg, L., Borssén, B., Olsson, T., and Gustafson, Y., 1998, “Fractures after Stroke,” Osteoporos Int, 8(1), pp. 92–95. DOI: 10.1007/s001980050053.

[9] Kapral, M. K., Fang, J., Alibhai, S. M. H., Cram, P., Cheung, A. M., Casaubon, L. K., Prager, M., Stamplecoski, M., Rashkovan, B., and Austin, P. C., 2017, “Risk of Fractures after Stroke,” Neurology, 88(1), pp. 57–64. DOI: 10.1212/WNL.0000000000003457.

[10] Longa, E. Z., Weinstein, P. R., Carlson, S., and Cummins, R., 1989, “Reversible Middle Cerebral Artery Occlusion without Craniectomy in Rats.,” Stroke, 20(1), pp. 84–91. DOI: 10.1161/01.STR.20.1.84.

[11] Peppen, R. P. V., Kwakkel, G., Wood-Dauphinee, S., Hendriks, H. J., Wees, P. J. V. der, and Dekker, J., 2004, “The Impact of Physical Therapy on Functional Outcomes after Stroke: What’s the Evidence?,” Clin Rehabil, 18(8), pp. 833–862. DOI: 10.1191/0269215504cr843oa.

[12] Bernhardt, J., English, C., Johnson, L., and Cumming, T. B., 2015, “Early Mobilization After Stroke Early Adoption but Limited Evidence,” Stroke, 46(4), pp. 1141–1146. DOI: 10.1161/STROKEAHA.114.007434.

[13] Borschmann, K., Iuliano, S., Ghasem-Zadeh, A., Churilov, L., Pang, M. Y. C., and Bernhardt, J., 2018, “Upright Activity and Higher Motor Function May Preserve Bone Mineral Density within 6 Months of Stroke: A Longitudinal Study,” Arch Osteoporos, 13(1), p. 5. DOI: 10.1007/s11657-017-0414-4.

[14] Park, S.-Y., Marasini, S., Kim, G.-H., Ku, T., Choi, C., Park, M.-Y., Kim, E.-H., Lee, Y.-D., Suh-Kim, H., and Kim, S.-S., 2014, “A Method for Generate a Mouse Model of Stroke: Evaluation of Parameters for Blood Flow, Behavior, and Survival,” Exp Neurobiol, 23(1), pp. 104–114. DOI: 10.5607/en.2014.23.1.104.

[15] Bouët, V., Freret, T., Toutain, J., Divoux, D., Boulouard, M., and Schumann-Bard, P., 2007, “Sensorimotor and Cognitive Deficits after Transient Middle Cerebral Artery Occlusion in the Mouse,” Experimental Neurology, 203(2), pp. 555–567. DOI: 10.1016/j.expneurol.2006.09.006.

[16] Bederson, J. B., Pitts, L. H., Tsuji, M., Nishimura, M. C., Davis, R. L., and Bartkowski, H., 1986, “Rat Middle Cerebral Artery Occlusion: Evaluation of the Model and Development of a Neurologic Examination,” Stroke, 17(3), pp. 472–476.

[17] Reglődi, D., Tamás, A., and Lengvári, I., 2003, “Examination of Sensorimotor Performance Following Middle Cerebral Artery Occlusion in Rats,” Brain Research Bulletin, 59(6), pp. 459–466. DOI: 10.1016/S0361-9230(02)00962-0.

[18] Prasad, J., Wiater, B. P., Nork, S. E., Bain, S. D., and Gross, T. S., 2010, “Characterizing Gait Induced Normal Strains in a Murine Tibia Cortical Bone Defect Model,” J. Biomech., 43(14), pp. 2765–2770. DOI: 10.1016/j.jbiomech.2010.06.030.

[19] Hurwitz, D. E., Sumner, D. R., Andriacchi, T. P., and Sugar, D. A., 1998, “Dynamic Knee Loads during Gait Predict Proximal Tibial Bone Distribution,” J. Biomech., 31(5), pp. 423–430. DOI: 10.1016/S0021-9290(98)00028-1.

[20] Frost, H. M., 2003, “Bone’s Mechanostat: A 2003 Update,” Anat. Rec., 275A(2), pp. 1081–1101. DOI: 10.1002/ar.a.10119.

[21] Ellman, R., Spatz, J., Cloutier, A., Palme, R., Christiansen, B. A., and Bouxsein, M. L., 2013, “Partial Reductions in Mechanical Loading Yield Proportional Changes in Bone Density, Bone Architecture, and Muscle Mass,” J. Bone. Miner. Res., 28(4), pp. 875–885. DOI: 10.1002/jbmr.1814.

[22] Leblond, H., L’Espérance, M., Orsal, D., and Rossignol, S., 2003, “Treadmill Locomotion in the Intact and Spinal Mouse,” J. Neurosci., 23(36), pp. 11411–11419.

[23] Kloos, A. D., Fisher, L. C., Detloff, M. R., Hassenzahl, D. L., and Basso, D. M., 2005, “Stepwise Motor and All-or-None Sensory Recovery Is Associated with Nonlinear Sparing after Incremental Spinal Cord Injury in Rats,” Exp. Neurol., 191(2), pp. 251–265. DOI: 10.1016/j.expneurol.2004.09.016.

[24] Bouxsein, M. L., Boyd, S. K., Christiansen, B. A., Guldberg, R. E., Jepsen, K. J., and Müller, R., 2010, “Guidelines for Assessment of Bone Microstructure in Rodents Using Micro–Computed Tomography,” J Bone Miner Res, 25(7), pp. 1468–1486. DOI: 10.1002/jbmr.141.

[25] Smith, L., Bigelow, E. M. R., and Jepsen, K. J., 2013, “Systematic Evaluation of Skeletal Mechanical Function,” Curr Protoc Mouse Biol, 3, pp. 39–67. DOI: 10.1002/9780470942390.mo130027.

[26] Dempster, D. W., Compston, J. E., Drezner, M. K., Glorieux, F. H., Kanis, J. A., Malluche, H., Meunier, P. J., Ott, S. M., Recker, R. R., and Parfitt, A. M., 2013, “Standardized Nomenclature, Symbols, and Units for Bone Histomorphometry: A 2012 Update of the Report of the ASBMR Histomorphometry Nomenclature Committee,” J Bone Miner Res, 28(1), pp. 2–17. DOI: 10.1002/jbmr.1805.

[27] Egan, K. P., Brennan, T. A., and Pignolo, R. J., 2012, “Bone Histomorphometry Using Free and Commonly Available Software,” Histopathology, 61(6), pp. 1168–1173. DOI: 10.1111/j.1365-2559.2012.04333.x.

[28] Schindelin, J., Arganda-Carreras, I., Frise, E., Kaynig, V., Longair, M., Pietzsch, T., Preibisch, S., Rueden, C., Saalfeld, S., Schmid, B., Tinevez, J.-Y., White, D. J., Hartenstein, V., Eliceiri, K., Tomancak, P., and Cardona, A., 2012, “Fiji: An Open-Source Platform for Biological-Image Analysis,” Nature Methods, 9(7), pp. 676–682. DOI: 10.1038/nmeth.2019.

[29] Morris, M. D., and Mandair, G. S., 2011, “Raman Assessment of Bone Quality,” Clin Orthop Relat Res, 469(8), pp. 2160–2169. DOI: 10.1007/s11999-010-1692-y.

[30] Gamsjaeger, S., Masic, A., Roschger, P., Kazanci, M., Dunlop, J. W. C., Klaushofer, K., Paschalis, E. P., and Fratzl, P., 2010, “Cortical Bone Composition and Orientation as a Function of Animal and Tissue Age in Mice by Raman Spectroscopy,” Bone, 47(2), pp. 392–399. DOI: 10.1016/j.bone.2010.04.608.

[31] Akkus, O., Adar, F., and Schaffler, M. B., 2004, “Age-Related Changes in Physicochemical Properties of Mineral Crystals Are Related to Impaired Mechanical Function of Cortical Bone,” Bone, 34(3), pp. 443–453. DOI: 10.1016/j.bone.2003.11.003.

[32] Borschmann, K. N., Rewell, S. S., Iuliano, S., Ghasem-Zadeh, A., Davey, R. A., Ho, H., Skeers, P. N., Bernhardt, J., and Howells, D. W., 2017, “Reduced Bone Formation Markers, and Altered Trabecular and Cortical Bone Mineral Densities of Non-Paretic Femurs Observed in Rats with Ischemic Stroke: A Randomized Controlled Pilot Study,” PLoS One; San Francisco, 12(3), p. e0172889. DOI: http://dx.doi.org.prox.lib.ncsu.edu/10.1371/journal.pone.0172889.

[33] Berman, A. G., Hinton, M. J., and Wallace, J. M., 2019, “Treadmill Running and Targeted Tibial Loading Differentially Improve Bone Mass in Mice,” Bone Rep, 10. DOI: 10.1016/j.bonr.2019.100195.

[34] Borschmann, K., Pang, M. Y. C., Bernhardt, J., and Iuliano-Burns, S., 2012, “Stepping towards Prevention of Bone Loss after Stroke: A Systematic Review of the Skeletal Effects of Physical Activity after Stroke,” International Journal of Stroke, 7(4), pp. 330–335. DOI: 10.1111/j.1747-4949.2011.00645.x.

[35] Gertz, K., Priller, J., Kronenberg, G., Fink, K. B., Winter, B., Schröck, H., Ji, S., Milosevic, M., Harms, C., Böhm, M., Dirnagl, U., Laufs, U., and Endres, M., 2006, “Physical Activity Improves Long-Term Stroke Outcome via Endothelial Nitric Oxide Synthase–Dependent Augmentation of Neovascularization and Cerebral Blood Flow,” Circ. Res., 99(10), pp. 1132–1140. DOI: 10.1161/01.RES.0000250175.14861.77.

[36] Somerville, J. M., Aspden, R. M., Armour, K. E., Armour, K. J., and Reid, D. M., 2004, “Growth of C57BL/6 Mice and the Material and Mechanical Properties of Cortical Bone from the Tibia,” Calcif. Tissue Int., 74(5), pp. 469–475. DOI: 10.1007/s00223-003-0101-x.

[37] Hughes, P. C., and Tanner, J. M., 1970, “The Assessment of Skeletal Maturity in the Growing Rat.,” J Anat, 106(Pt 2), pp. 371–402.

[38] Horton, J. A., Bariteau, J. T., Loomis, R. M., Strauss, J. A., and Damron, T. A., 2008, “Ontogeny of Skeletal Maturation in the Juvenile Rat,” The Anatomical Record, 291(3), pp. 283–292. DOI: 10.1002/ar.20650.

[39] van der Meulen, M. C. H., Jepsen, K. J., and Mikić, B., 2001, “Understanding Bone Strength: Size Isn’t Everything,” Bone, 29(2), pp. 101–104. DOI: 10.1016/S8756-3282(01)00491-4.

[40] Kohn, D. H., Sahar, N. D., Wallace, J. M., Golcuk, K., and Morris, M. D., 2008, “Exercise Alters Mineral and Matrix Composition in the Absence of Adding New Bone,” Cells Tissues Organs; Basel, 189(1–4), pp. 33–7.

[41] Gardinier, J. D., Rostami, N., Juliano, L., and Zhang, C., 2018, “Bone Adaptation in Response to Treadmill Exercise in Young and Adult Mice,” Bone Rep, 8, pp. 29–37. DOI: 10.1016/j.bonr.2018.01.003.

[42] Wallace, J. M., Rajachar, R. M., Allen, M. R., Bloomfield, S. A., Robey, P. G., Young, M. F., and Kohn, D. H., 2007, “Exercise-Induced Changes in the Cortical Bone of Growing Mice Are Bone- and Gender-Specific,” Bone, 40(4), pp. 1120–1127. DOI: 10.1016/j.bone.2006.12.002.

[43] Fassbender, K., Rossol, S., Kammer, T., Daffertshofer, M., Wirth, S., Dollman, M., and Hennerici, M., 1994, “Proinflammatory Cytokines in Serum of Patients with Acute Cerebral Ischemia: Kinetics of Secretion and Relation to the Extent of Brain Damage and Outcome of Disease,” Journal of the neurological sciences, 122(2), pp. 135–139.

[44] Tuttolomondo, A., Di Raimondo, D., di Sciacca, R., Pinto, A., and Licata, G., 2008, “Inflammatory Cytokines in Acute Ischemic Stroke,” Current Pharmaceutical Design; Schiphol, 14(33), pp. 3574–89. DOI:http://dx.doi.org.prox.lib.ncsu.edu/10.2174/138161208786848739.

[45] Pang, M. Y. C., Yang, F. Z. H., and Jones, A. Y. M., 2013, “Vascular Elasticity and Grip Strength Are Associated With Bone Health of the Hemiparetic Radius in People With Chronic Stroke: Implications for Rehabilitation,” PHYS THER, 93(6), pp. 774–785. DOI: 10.2522/ptj.20120378.

[46] Stegen, S., and Carmeliet, G., 2018, “The Skeletal Vascular System – Breathing Life into Bone Tissue,” Bone, 115, pp. 50–58. DOI: 10.1016/j.bone.2017.08.022.

[47] Wong, I. P. L., Zengin, A., Herzog, H., and Baldock, P. A., 2008, “Central Regulation of Bone Mass,” Seminars in Cell & Developmental Biology, 19(5), pp. 452–458. DOI: 10.1016/j.semcdb.2008.08.001.

